# Loss of the mitochondrial redox gene *GLRX2* underlies somersaulting behavior in Darwin’s pigeons

**DOI:** 10.64898/2026.11.12.766293

**Authors:** Si Si, Ling-Yan Su, Lan Long, You Qian, Wen-Di Ma, Xin Gan, Kun Jiang, Guo-Hao Wang, Zoltán Bagi, Daniele Bigi, Hamed Kharrati-Koopaee, Guang-Ze Zheng, Li Ma, Zhuang Wu, Ming-Li Li, Hong-Man Chen, Qiu Pan, Wen-Jing Peng, Zhu Bu, Wen Gao, Jian Li, Ci-Rong Liu, Jia Pan, Jiu-Lin Du, Zhen Liu, Chang Zhang, Chen Yan, Yong-Gang Yao, Lihong Gu, Li-Zhi Lu, Jun Lu, Kusza Szilvia, Ali Esmailizadeh, Guojie Zhang, Ming-Shan Wang

## Abstract

Darwin described the somersaulting behavior of pigeons as “one of the most remarkable inherited habits ever recorded”, yet its molecular basis remained unknown. We show that somersaulting pigeon brains exhibit mitochondrial redox dysfunction and identify an 11-bp frameshifting deletion in *GLRX2*, a mitochondrial redox enzyme, that abolishes protein function and is highly-enriched in two independently selected breeds. *GLRX2* deficiency in mice and zebrafish recapitulates mitochondrial dysfunction, oxidative stress and impaired motor coordination seen in pigeons. Single-nucleus transcriptomics identifies cerebellar Golgi and Purkinje cells as selectively vulnerable, with disrupted inhibitory signaling and synaptic remodeling. These findings demonstrate how relaxation of purifying selection during domestication can expose a deleterious mitochondrial variant to artificial selection, allowing gene loss to serve as a substrate for behavioral evolution.

## Main Text

Understanding how complex behaviors arise from genetic variation remains a central challenge in biology. Animal domestication offers a powerful framework with which to address this question (*1–3*), since artificial selection can rapidly amplify unusual, even maladaptive, phenotypes that would not persist in a nature (*4*). The domestic pigeon (*Columba livia*) is a classic case. Charles Darwin was fascinated by the extensive phenotypic variation among pigeon breeds and used them to illustrate how sustained selection acting on a single lineage can generate forms as divergent as those seen among distinct genera (*5, 6*). Since then, “Darwin’s pigeons” have served as a key model for studying the genetic architecture of rapid phenotypic evolution and the diversification of complex traits (*5, 7, 8*).

Derived from the Rock pigeon and domesticated in Neolithic times (*9*), domestic pigeons now encompass more than 350 breeds exhibiting extensive and extreme variations in morphology and behavior. Among these, somersaulting (or tumbling) behavior is one of the most unusual traits. Somersaulting pigeons perform involuntary backward flips or continuous rolling movements during flight or on the ground, either spontaneously or in response to external stimuli (*10*) (**Fig. 1A and movies S1 to S3**). This behavior has been documented since at least the sixteenth century (*11*), and Darwin described it as “one of the most remarkable inherited habits or instincts ever recorded” (*5*). In natural conditions, such loss of balance and coordination would likely be lethal through increased predation risk. Yet under artificial selection, this seemingly maladaptive motor pattern has persisted and diversified into a wide spectrum of forms and severities, from flying tumblers that perform aerial somersaults to flightless Parlor rollers that repeatedly roll along the ground (**Fig. 1A and movies S1 to S3**).

**Fig. 1.**
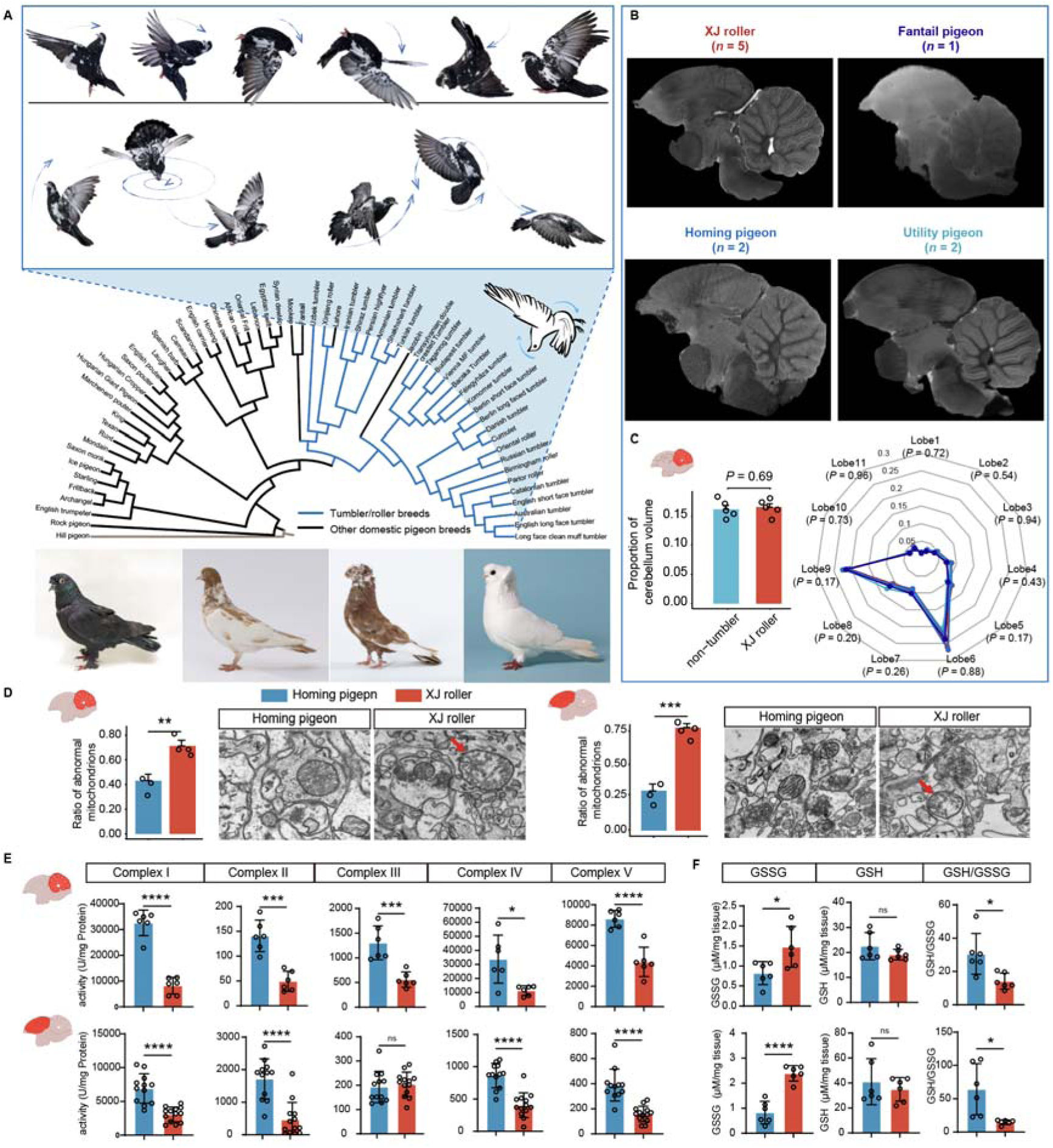
Somersaulting pigeons exhibit mitochondrial ultrastructural and functional defects without overt macroscopic brain abnormalities. (A) Schematic illustration of somersaulting behavior, a neighbor-joining tree of domestic pigeon breeds based on genome wide variations, and images of four representative tumbler and roller breeds spanning the behavioral spectrum of the trait. From left to right, the breeds shown are Xinjiang roller, Parlor roller, Transylvanian double-crested tumbler, and Félegyháza tumbler. Branches corresponding to tumbler and roller breeds are shown in blue. (B) Representative sagittal MRI sections from Xinjiang roller, fantail pigeon, homing pigeon, and utility pigeon. No overt gross anatomical abnormalities were detected in somersaulting pigeon brains. **(C)** MRI-based quantification of relative cerebellum volume and cerebellar lobule organization. Left, cerebellum volume normalized to total brain volume in somersaulting (Xinjiang roller) and non-somersaulting pigeons. Right, proportional volumes of individual cerebellar lobules normalized to total cerebellum volume. *P* values shown for each lobule indicate comparisons of normalized lobular volume between somersaulting and non-somersaulting pigeons. **(D)** Transmission electron microscopy of neuronal synaptic regions in the cerebellum (left) and cerebrum (right) of homing pigeons and Xinjiang rollers showing the mitochondrial morphology. Red arrows mark abnormal mitochondria. Quantification of abnormal mitochondria for each tissue is shown at left of each image set. Scale bars, 200 nm. **(E)** Activities of mitochondrial respiratory chain complexes I to V in the cerebellum (top) and cerebrum (bottom). **(F)** GSSG, GSH, and GSH/GSSG ratios in the cerebellum (top) and cerebrum (bottom). Somersaulting pigeons (blue bars) showed frequent dysmorphic mitochondria, reduced respiratory chain activity, increased GSSG, and decreased GSH/GSSG ratios compared with homing pigeons (red bars). Bars indicate mean ± s.e.m.; points indicate individual animals or biological replicates. Statistical significance was assessed by unpaired two-tailed *t* test. ns, not significant; \**P* < 0.05, \*\**P* < 0.01, \*\*\**P* < 0.001, and \*\*\*\**P* < 0.0001

The evolutionary and genetic basis of somersaulting has been debated for over a century. Multiple mechanistic hypotheses have been proposed, including dysfunction of cerebellar or vestibular systems, altered spinal reflexes, abnormal neurotransmission, and neuromuscular defects (*12–16*), but anatomical and pharmacological investigations have failed to identify a consistent causal explanation. Breeding studies indicate that somersaulting is highly heritable and likely controlled by a major recessive locus (*ro*), possibly with additional modifiers, yet its molecular identity remains unknown (*17, 18*). Moreover, it is also unclear whether the diverse somersaulting phenotypes observed across breeds reflect a shared genetic mechanism or independent genetic solutions (*17*).

Here, we combined population genomics, multiscale neuroimaging, single-nucleus transcriptomics, and cross-species functional genetics to investigate the molecular basis of somersaulting behavior. Starting from ultrastructural and functional characterization of somersaulting pigeon brain, we trace the trait to an 11-bp loss-of-function deletion in *GLRX2*, a mitochondrial redox enzyme. We then characterize the physiological and transcriptional consequences of *GLRX2* loss across cell types and brain regions, and test its causal role in locomotor coordination through knockout experiments in mice and zebrafish.

## Result

### Somersaulting pigeons exhibit mitochondrial ultrastructural and functional defects in the brain

Early studies proposed cerebellum, the vestibular system or the spinal reflex circuits as candidate mechanisms for somersaulting behavior (*12, 15*), but detailed anatomical examinations of the brain, inner ear and musculature have failed to identify consistent structural correlates (*12, 17, 19*). These analyses, however, relied largely on gross anatomical observations and did not systematically examine brain ultrastructural. To address this gap, we compared the neuroanatomy and subcellular features of somersaulting pigeons (Xinjiang rollers) with non-somersaulting control pigeons at complementary macroscopic and ultrastructural scales (**Fig. 1, B to D**).

At the macroscopic level, we performed high-resolution whole-brain magnetic resonance imaging (MRI; 9.4 T Bruker system) and revealed no overt neuroanatomical abnormalities in somersaulting pigeons compared with non-somersaulting pigeons (*n* = 5 biological replicates per group; **Fig. 1, B and C**, **and fig. S1**). Three-dimensional reconstruction and volumetric segmentation showed that the relative volumes of cerebellum and cerebrum, as well as the proportional representation of individual cerebellar lobules, were broadly comparable across breeds, with no evidence of regional deformation or malformation (**Fig. 1, B and C, and fig. S1**). Somersaulting behavior is therefore unlikely to be caused by gross neuroanatomical abnormalities at this resolution.

At the ultrastructural level, transmission electron microscopy (TEM) revealed that neuronal synapses in somersaulting pigeons contained a high frequency of dysmorphic mitochondria, in both the cerebellum and the cerebrum (*n* = 3 biological replicates per group, **Fig. 1D**). These mitochondria exhibited pronounced swelling, vacuolisation, fragmented cristae, and severe disruption of the mitochondrial membrane (**Fig. 1D**), whereas neurons from homing pigeons displayed normal cellular architecture. The concentration of mitochondrial damage within synaptic regions suggests that the bioenergetic demands of synaptic transmission may render these compartments particularly vulnerable.

Consistent with the ultrastructural damage observed, mitochondrial respiratory function was impaired in somersaulting pigeon brains. Respiratory chain supercomplexes (RCS) activities were significantly reduced, with the cerebellum showing a marked decrease across complexes I– V **(Fig. 1E**). Because cristae morphology critically influences the assembly of RCS and overall bioenergetic efficiency (*20, 21*), these functional deficits likely reflect the structural compromise observed by TEM. To assess whether mitochondrial dysfunction was accompanied by redox imbalance, a phenomenon often co-occurring with mitochondrial dysfunction (*22*), we measured glutathione redox status, a sensitive indicator of cellular redox status and reactive oxygen species (ROS) accumulation (*23*). While reduced glutathione (GSH) levels remained comparable between somersaulting pigeons and controls (n = 6 biological replicates per group per tissue), oxidized glutathione (GSSG) was significantly elevated in the cerebrum (*P* < 0.0001) and cerebellum (*P* = 0.02), and GSH/GSSG ratio was also markedly decreased in both regions of somersaulting pigeons (*P* < 0.05 for each; **Fig. 1F**), implying a state of severe oxidative stress.

Together, these results reveal that the somersaulting pigeon brain is characterized by mitochondrial structural and functional compromise, with disrupted oxidative phosphorylation (OXPHOS) system and a shift toward a pro-oxidative cellular environment, in the absence of gross neuroanatomical abnormality. These findings implicate mitochondrial dysfunction as a key cellular feature of the somersaulting phenotype.

### Mitochondrial gene programs are dysregulated in somersaulting pigeon brains

The ultrastructural and biochemical defects identified above implicate mitochondrial dysfunction as a cellular feature of the somersaulting pigeon phenotype. To identify transcriptional signatures associated with somersaulting, we profiled gene expression across the cerebrum, cerebellum, and midbrian of Xinjiang rollers (somersaulting), homing pigeons, and utility pigeons (*n* = 6 biological replicates per breed) using a newly assembled chromosome-scale reference genome for the pigeon (**fig. S2**).

Weighted Gene Co-expression Network Analysis (WGCNA) (*24*) identified 22 co-expression modules, among which the MEdarkred module (containing 31 genes) exhibited the strongest correlation with somersaulting behavior (*r* = −0.46, *P* = 7×10^-4^) but no significant association with brain region (**Fig. 2A**), indicating that this module captures phenotype-related transcriptional variation rather than regional differences. Genes in this module were functionally enriched in mitochondrial dynamics (e.g., regulation of fission and cytochrome c release) and neurological development (e.g., neuroblast delamination, neural crest formation, forebrain differentiation) (**Fig. 2A**), and were significantly downregulated in the somersaulting group (**table S1**). Among them were several genes essential for mitochondrial integrity and bioenergetics, including the complex I subunit *NDUFA6* (25), the mitochondrial nucleotide kinase *CMPK2* (26), the innate immune regulator *SAMD9L* (*27*), mutations in which have been implicated in human spinocerebellar ataxia (*28*).

**Fig. 2.**
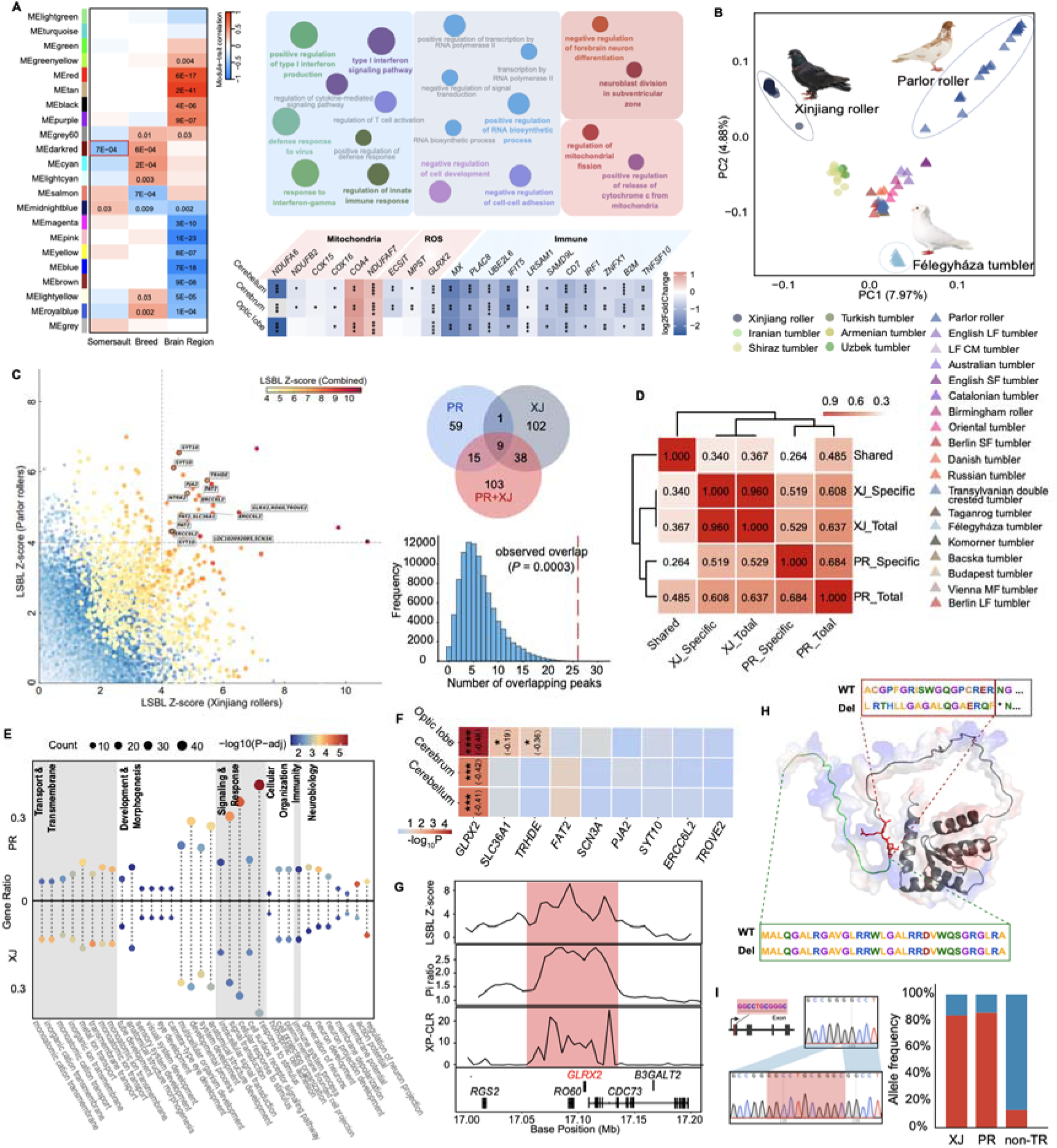
Multi-omic analyses identify *GLRX2* as a candidate gene underlying somersaulting behavior in pigeons. **(A)** WGCNA identified modules associated with somersaulting behavior and summarized their functional enrichments. The heat map below shows representative differentially expressed genes related to mitochondrial function, redox regulation, and immune pathways across brain regions. **(B)** PCA of domestic pigeon breeds based on whole-genome SNPs. Xinjiang roller, Parlor roller, and Félegyháza tumbler are differentiated from the remaining tumbler breeds. Félegyháza tumbler corresponds to a lineage in which the somersaulting phenotype has been lost. **(C)** LSBL scans in Xinjiang rollers (XJ), Parlor rollers (PR), and combined group (XJ+PR). Left, comparison of LSBL *Z*-scores in the three analyses, with point color indicating the LSBL *Z*-score from the combined-group analysis. Labeled points denote windows containing genes, unlabeled points denote windows in noncoding regions. Top right, overlap of genes showing signatures of selection. Bottom right, permutation test showing that the observed overlap is greater than expected by chance. **(D)** Semantic similarity analysis of functional enrichment terms for genes under selection. Shared, genes detected in XJ and PR; XJ_specific and PR_specific, genes unique to each analysis; XJ_total and PR_total, total genes detected in each analysis. **(E)** Functional enrichment of genes under selection in XJ and PR. **(F)** Expression of candidate genes shared among three selection scans across brain tissues. Asterisks denote significant differential expression, and values in parentheses indicate log_2_Fold Change. **(G)** Selection signals across the *GLRX2* locus, including LSBL *Z*-score, nucleotide diversity ratio, and XP-CLR. The highlighted interval marks the candidate selected region encompassing *GLRX2*. **(H)** Predicted GLRX2 protein structure and the structural consequence of the 11-bp deletion. Green ribbons denote conserved regions of the WT protein, red ribbons denote the frameshift-derived peptide, and gray ribbons denote the truncated portion downstream of the premature stop codon introduced by the deletion. (I) Sequence and genomic location of the *GLRX2* 11-bp deletion (left) and allele frequencies of the deletion in Xinjiang roller (XJ), Parlor roller (PR), and non-tumbler/roller (non-TR) pigeons (right). The blue bar denotes the WT allele and red denotes the deletion allele.

Differentially expressed genes (DEGs) analysis further supports this finding. DEGs were functionally enriched in the regulation of mitochondrial membrane potential and depolarization (**table S2**). Notably, the mitochondrial complex I accessory subunits *NDUFA6* and *NDUFB2* (*29*), the cytochrome c oxidase assembly factors *COX15* (*30*) and *COX16* (*31*), the complex I assembly and mitochondrial ROS regulator *ECSIT* (*32, 33*), the mitochondrial glutaredoxin and redox sensor *GLRX2* (*34*), and the sulfur-metabolizing enzyme *MPST* (*35*), were significantly downregulated in somersaulting pigeons (**Fig. 2A**), consistent with the ultrastructural mitochondrial defects and redox imbalance observed at the biochemical level (**Fig. 1, D to F**). The overall transcriptional signature combined respiratory chain and antioxidant suppression, parallels patterns described in human mitochondrial ataxias (*36–38*).

Together, these transcriptomic signatures confirmed that persistent bioenergetic stress and impaired redox buffering are features of the somersaulting pigeon brains, and point to a genetically encoded disruption of mitochondrial function underlying the observed cellular pathology.

### A loss-of-function deletion in *GLRX2* is associated with somersaulting behavior

To investigate the genetic basis of the transcriptional and mitochondrial phenotypes described above, we performed whole-genome sequencing (WGS) of 171 pigeons from nine breeds at ∼27× coverage for each (**table S3**), and integrated these data with a global panel of 168 publicly available genomes, yielding more than 30 million high-quality variants across 339 individuals (**table S3**). Population clustering, principal component analysis, and structure analyses revealed that the 28 tumbling and rolling breeds (*n* = 178) cluster together and share a common ancestry (**Fig. 1A and fig. S3A**), while exhibiting pronounced genetic differentiation among lineages, with Parlor rollers (PR) and Xinjiang rollers (XJ) showing particularly strong divergence from other somersaulting breeds (**Fig. 2B and fig. S3, B and C**). These results support a single ancestral origin for tumbler and roller pigeons, followed by diversification under lineage-specific selection for distinct performance and morphological traits (*5, 39*).

Because somersaulting is likely controlled by a major recessive locus with additional modifiers (*17*), we performed locus-specific branch-length (LSBL) analyses to identify lineage-specific selection signals, using PR and XJ as focal populations and non-performing tumbler breeds and non-tumbler breeds as outgroups. We identified 169 protein-coding genes with signature of selection (hereafter PSGs) in the XJ and 92 in the PR breed (*Z* > 4.0; **table S4**). Although outlier-based approach can be susceptible to false positives from demographic history and genetic drift (*40*), the convergence of independent signals across two independently bred lineages mitigates this concern. Ten genes (*SCN3A, SYT10, GLRX2, FAT2, PJA2, SLC36A, TRHDE, ERCC6L2*, *TROVE2,* and *NTRK2*) were shared between the two lineages (**Fig. 2C**), an overlap significantly greater than would be expected by chance (*P* = 0.0003; **Fig. 2C**). Despite limited gene-level overlap, PSGs from the two lineages converged on similar biological pathways (semantic correlation >0.50; **Fig. 2D**), including neuronal and axonal development, synaptic signaling, neuronal action potentials, motor behavior and immune system processes (**Fig. 2E**). Several mitochondrial function genes (*GLRX2, NNT, HINT2* and *SLC25A39*) also exhibited strong selection signals (*Z* > 4.0, at least in XJ or PR; **table S4**), implicating mitochondrial pathways as shared targets of selection across somersaulting lineages.

Combined LSBL analysis of XJ and PR together recapitulated strong selection signals for nine of the ten shared genes (*Z* > 4.0; **Fig. 2C and table S4**), and genome-wide *F_ST_* scanning confirmed that seven of the ten (*SYT10, GLRX2, FAT2, PJA2, TRHDE, ERCC6L2*, and *TROVE2*) fell above the 99.9th percentile (**fig. S4**). To identify which of these candidates might causally contribute to the phenotype, we examined their expression across brain regions. *GLRX2* emerged as the most compelling locus, which was the only shared PSG consistently and significantly downregulated across all three brain regions in somersaulting pigeons relative to controls (**Fig. 2F**). Independent selection metrics further supported this candidacy, including elevated cross-population composite likelihood ratio (XP-CLR) scores, increased π-ratio, and high LSBL (*Z* = 9.24; in combined group) (**Fig. 2G and table S4**).

*GLRX2* (glutaredoxin 2) encodes a mitochondrial thiol-disulfide oxidoreductase central to bioenergetic function and redox homeostasis (*41*). Its deficiency has been linked to impaired respiratory chain efficiency (*42*) and to multiple pathological conditions, including cardiac hypertrophy, impaired angiogenesis, cataract formation and neurological disorders (*43–46*), to our knowledge, a role in motor behavior has not been widely reported. Screening for causal variation within *GLRX2* revealed an 11-bp deletion in exon 1 (**Fig. 2, H and I**) that introduces a frameshift after amino acid 34, generating a premature stop codon at position 52 and eliminating 127 downstream amino acids (**Fig. 2H**), predicted to abolish *GLRX2* function. Because premature termination codons trigger nonsense-mediated mRNA decay (*47*), the reduced *GLRX2* expression observed in the somersaulting pigeon brain and confirmed by quantitative PCR (**fig. S5A**) is likely a consequence of this mutation. To determine whether any full-length transcripts initiating downstream of the first exon or skipping the deletion were present, we assembled and analyzed the transcriptomes. However, no such transcripts were detected (**fig. S5B**), indicating that the 11-bp deletion in exon 1 likely disrupts gene function, leading to the complete pseudogenization of *GLRX2*.

The deletion allele was highly enriched in the two somersaulting breeds, PR (86.36%) and XJ (84.21%), compared with 13.45% in other breeds (Fisher’s exact test, *P* = 1.17×10^−46^; **Fig. 2I**). Consistent with previous studies showed that somersaulting is a recessive trait (*17*), the deletion was predominantly homozygous in PR (77.27%) and XJ (73.68%), but was rarely observed in other breeds (5.26%; **fig. S6A**). Genotyping of an independent cohort of 41 Xinjiang rollers confirmed this pattern, with an allele frequency of 87.80% and 32 of 41 individuals being homozygotes (**fig. S6B**). The consistency between the initial and replication cohorts, both exhibiting allele frequencies approaching 90%, suggests that this allele is approaching fixation. This finding effectively precludes the possibility of sampling bias as an explanation for the observed association. Notably, fantail pigeons, a sister lineage **(Fig. 1A and fig. S6A**) to the Xinjiang roller that frequently exhibits spasm-like backward toppling when excited (*12*), which is reminiscent of the abnormal tail posture and loss of balance observed during pigeon somersaulting (*10*). We found that fantail pigeons also harbor the homozygous deletion at a high frequency (63.64%; **fig. S6A**), extending the genotype-phenotype association beyond the core somersaulting breeds.

### *GLRX2* loss recapitulates locomotor coordination defects in model organisms

To test whether *GLRX2* loss is sufficient to impair locomotor coordination, we used zebrafish and mice as model systems, given the current challenges of gene editing in avian species and the broad conservation of GLRX2 across vertebrates (**fig. S7**). Examination of chickenGTEx data (https://ngdc.cncb.ac.cn/chickengtex/) showed that *GLRX2* is broadly expressed in various tissues (**fig. S8**), which is consistent with its core mitochondrial role. We applied two GLRX2-knockout (KO) zebrafish lines using CRISPR/Cas9-mediated genome editing (**fig. S9**) and a GLRX2-KO mouse line generated by homologous recombination (**fig. S10**) to assess motor behavior in both species.

In zebrafish, we examined larval locomotor behavior following hydrodynamic stimulation at 72 hours post-fertilization (hpf) (**Fig. 3A**). To quantify swimming patterns, we developed a tracking method using the Lattice Boltzmann Model (LBM) to reconstruct three-dimensional trajectories. Wild-type (WT) larvae exhibited the typical high-velocity escape response with linear or curvilinear trajectories (**Fig. 3B and movie S4**). In contrast, GLRX2-KO larvae displayed severely uncoordinated swimming characterized by repetitive circular motion (**Fig. 3B and movie S5**). GLRX2-KO larvae from two independent lines showed a robust increase in rotational swimming across multiple independent batches compared with WT controls (**Fig. 3C**). Within the same developmental cohort, individuals from KO genotypes exhibited circling frequencies of 56.16%–84.21%, whereas circling occurring in only 0%–9.84% of WT individuals (**Fig. 3C**). This pronounced and reproducible phenotype, reminiscent of the rotational behavior observed in somersaulting pigeons, indicates that *GLRX2* loss produces a predictable deficit in locomotor coordination.

**Fig. 3.**
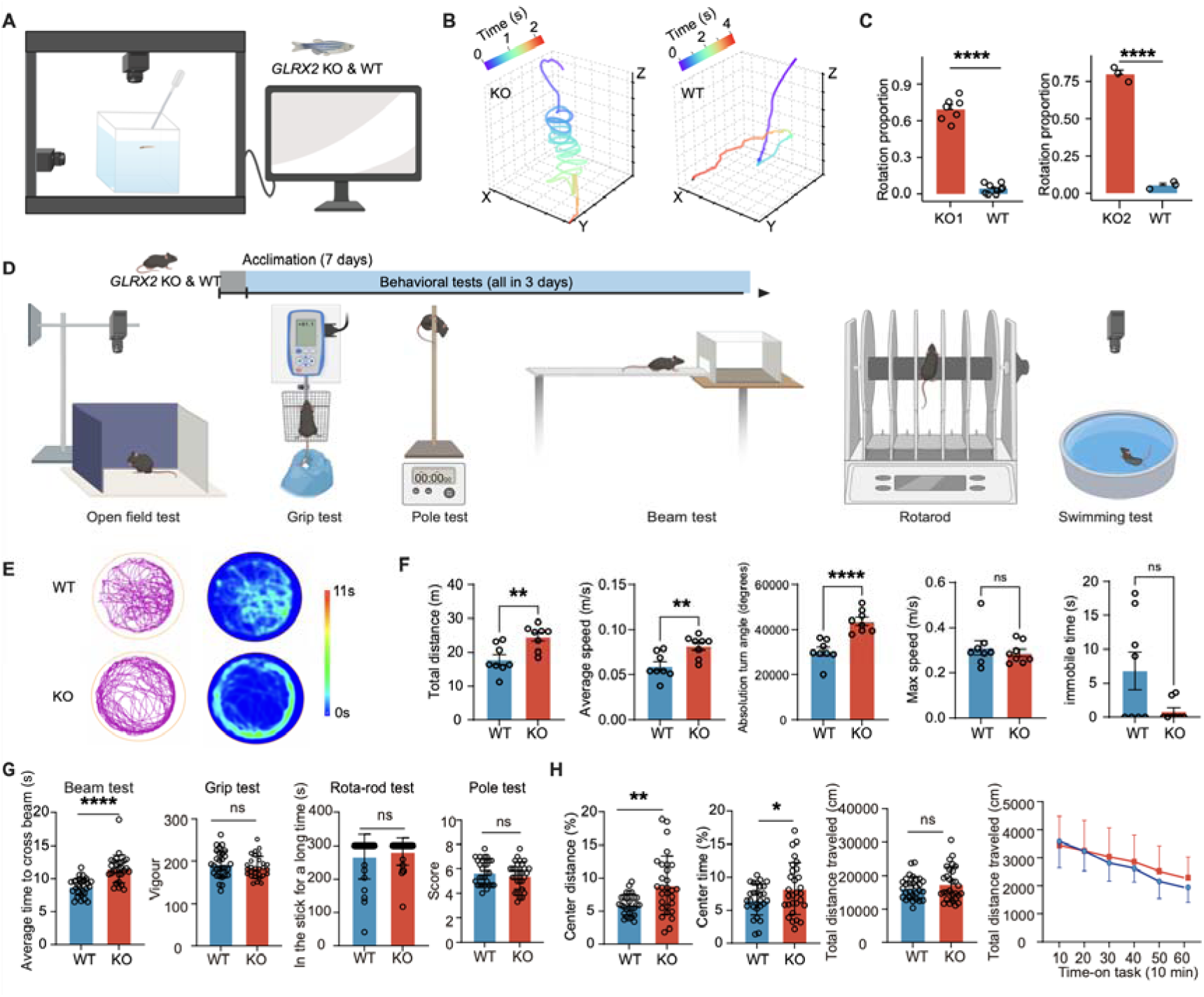
*GLRX2* deficiency causes locomotor coordination defects in zebrafish and mice. **(A)** Experimental setup for behavioral analysis of zebrafish larvae. GLRX2-KO and WT larvae were subjected to hydrodynamic stimulation, and three-dimensional locomotor trajectories were recorded using a multi-view tracking system. **(B)** Representative three-dimensional swimming trajectories of *GLRX2*-KO and WT zebrafish larvae. Color gradients indicate elapsed time. **(C)** Quantification of rotational swimming behavior in two independent GLRX2-KO zebrafish lines (KO1 and KO2) and WT controls. The proportion of larvae exhibiting rotational swimming is shown for each genotype. **(D)** Experimental design for behavioral testing of GLRX2-KO and WT mice. **(E)** Representative swimming trajectories and residence-time maps of WT and GLRX2-KO mice during the swimming test. **(F)** Quantification of swimming behavior, including total distance, average speed, absolute turn angle, maximum speed, and immobile time. **(G)** Quantification of motor coordination and strength in GLRX2-KO and WT mice using balance beam, grip strength, rotarod, and pole tests. **(H)** Open-field test measurements showing center distance, center time, and total distance traveled over time in GLRX2-KO and WT mice. Bars indicate mean ± s.e.m.; points indicate individual animals. Statistical significance was assessed by unpaired two-tailed *t* test. ns, not significant; \**P* < 0.05, \*\**P* < 0.01, \*\*\**P* < 0.001, and \*\*\*\**P* < 0.0001.

In mice, GLRX2-KO animals showed a comparable coordination phenotype (**Fig. 3D**). In an open-water swimming test, GLRX2-KO mice displayed a pronounced rotational bias characterized by sustained unilateral circling, whereas WT controls exhibited irregular exploratory trajectories (**Fig. 3, E and F**). Consistent with impaired balance and coordination, GLRX2-KO mice also required significantly more time to traverse the balance beam (**Fig. 3G and fig. S11**). By contrast, grip strength, rotarod performance, and pole-climbing ability were not significantly affected (**Fig. 3G and fig. S11**), suggesting that the deficit primarily reflects impaired coordinative control rather than reduced muscular strength or general motor endurance. In addition, GLRX2-KO mice spent more time and travelled greater distances in the center of the open field (**Fig. 3H**). The basis of this phenotype is uncertain and may reflect either reduced thigmotaxis or disrupted spatial-orientation control secondary to the coordination deficit.

Collectively, the consistent impairment of balance and locomotor coordination across zebrafish, mice, paralleling the phenotype observed in somersaulting pigeons, supports a conserved role for *GLRX2* in motor coordination and indicates that its loss is sufficient to disrupt the neural mechanisms underlying locomotor control.

### *GLRX2* deficiency induces mitochondrial dysfunction and neuropathology

Having established that *GLRX2* loss impairs locomotor coordination in zebrafish and mice, we next asked whether it also recapitulate the mitochondrial pathology observed in somersaulting pigeon brains (**Fig. 1, D to F**). TEM revealed that neuronal mitochondria in both GLRX2-KO mice and zebrafish displayed prominent structural abnormalities, including swelling, vacuolization, cristae disruption, and membrane damage (**Fig. 4A**). Damaged mitochondria were frequently engulfed by autophagosomes in brain tissues (**Fig. 4B**), indicating elevated mitophagy. These abnormalities closely paralleled those observed in somersaulting pigeon brains, supporting *GLRX2* loss as their cause.

**Fig. 4.**
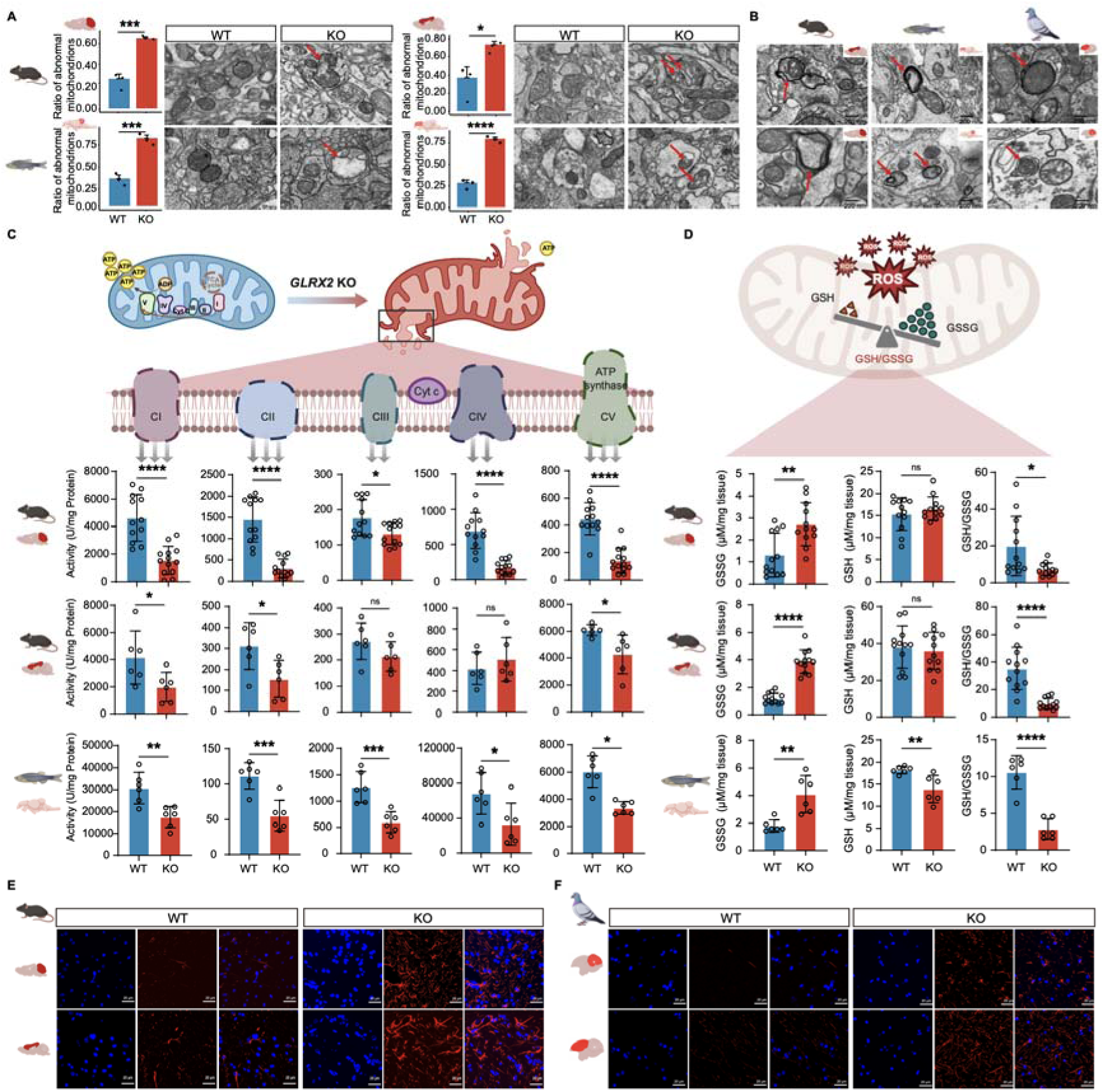
*GLRX2* deficiency induces conserved mitochondrial and neuropathological abnormalities. **(A)** Transmission electron microscopy of neuronal mitochondria in mouse and zebrafish brain tissues. Red arrows indicate abnormal mitochondria. Bar plots show the proportion of abnormal mitochondria. **(B)** Representative transmission electron micrographs showing damaged mitochondria enclosed within autophagic vacuoles in mouse, zebrafish, and pigeon brains. Red arrows indicate mitochondria undergoing mitophagy. **(C)** Schematic of mitochondrial respiratory chain dysfunction after *GLRX2* loss and quantification of respiratory chain complex activities in mouse and zebrafish brain tissues. **(D)** Schematic of redox imbalance and measurements of oxidized glutathione (GSSG), reduced glutathione (GSH), and the GSH/GSSG ratio in mouse and zebrafish brain tissues. **(E)** GFAP immunostaining in the cerebrum and cerebellum of WT and GLRX2-KO mice. **(F)** GFAP immunostaining in the cerebrum and cerebellum of homing and tumbler pigeons. Bars indicate mean ± s.e.m.; points indicate individual animals or biological replicates. Statistical significance was assessed by unpaired two-tailed *t* test. ns, not significant; \**P* < 0.05, \*\**P* < 0.01, \*\*\**P* < 0.001, and \*\*\*\**P* < 0.0001.

Consistent with the ultrastructural damage, mitochondrial respiratory function was impaired in GLRX2-KO animals (**Fig. 4C**). RCS activity was broadly reduced, with significant suppression of complexes I, II, and V in both mice and zebrafish (**Fig. 4C**). Oxidized glutathione (GSSG) was elevated while reduced glutathione remained largely unchanged, resulting in a reduced GSH/GSSG ratio across examined brain tissues (**Fig. 4D**). This pattern mirrors the bioenergetic and redox compromise observed in somersaulting pigeon brains (**Fig. 1, E and F**), indicating a conserved disruption of mitochondrial OXPHOS and redox homeostasis from *GLRX2* loss.

Because excessive ROS accumulation can promote neuronal damage through neuroinflammatory processes, we next evaluated astrocyte activation by immunofluorescence staining for glial fibrillary acidic protein (GFAP) (*48*). GFAP immunoreactivity was markedly increased in both the cerebellum and cerebrum of GLRX2-KO mice, accompanied by hypertrophic astrocytic morphology consistent with astrogliosis (**Fig. 4E**). Somersaulting pigeons exhibited the same elevated GFAP immunoreactivity in both brain homologous regions (**Fig. 4F**). This shared neuroinflammatory signature indicates that *GLRX2* deficiency drives the glial activation component of the phenotype across species. To determine whether this inflammatory state was accompanied by overt cell death, we performed TUNEL staining in both species. However, no apparent apoptotic signals were detected (**fig. S12**), suggesting that the pathology remains sublethal.

These results establish *GLRX2* deficiency recapitulates the full spectrum of mitochondrial, redox, and neuroinflammatory pathology observed in somersaulting pigeon brains, without triggering overt neuronal apoptosis. Together with the locomotor coordination deficits described above, these findings causally implicate *GLRX2* loss in the somersaulting phenotype.

### Golgi and Purkinje cells are selectively vulnerable in somersaulting pigeons

The above results established that *GLRX2* deficiency compromises mitochondrial integrity and impairs locomotor coordination, but it remains unclear whether *GLRX2* deficiency alters cell type–specific transcriptional programs in the brain, and how such changes might contribute to the somersaulting phenotype. To address this, we performed single-nuclei RNA sequencing (snRNA-seq) data on the cerebellum and cerebrum of Xinjiang rollers (*n* = 3) and homing pigeons (*n* = 3; see details in Methods). After quality control, we obtained 54,460 cerebellar and 64,876 cerebral nuclei (median UMI counts per cell = 3,918 and median detected genes per cell = 1,695; **table S5 and figs. S13 and S14**), and clustering identified 16 cerebellar and 11 cerebral cell populations encompassing the major neuronal and glial cell types in each region (**Fig. 5A and fig. S15**), based on established cell marker genes (*49, 50*).

**Fig. 5.**
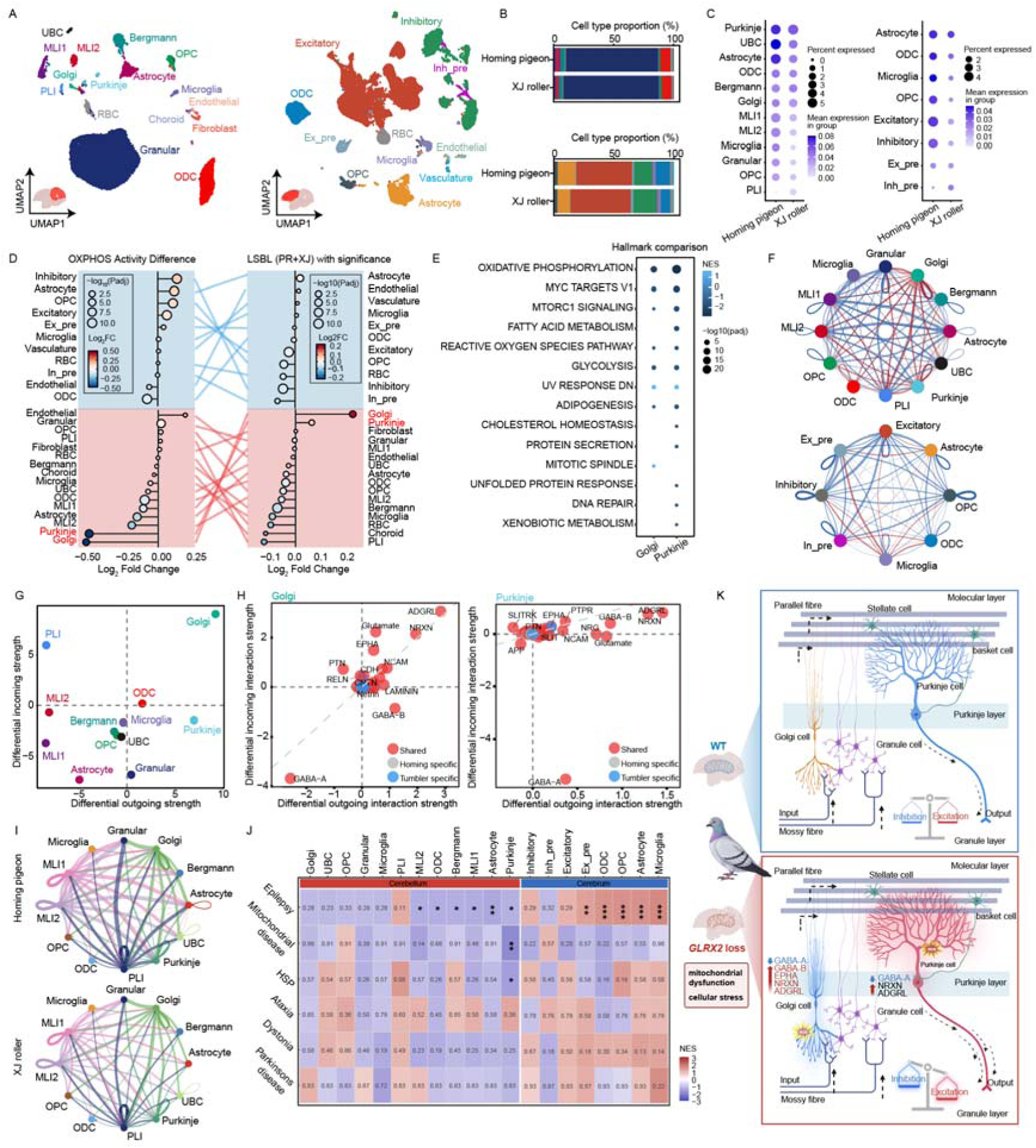
snRNA-seq identifies selective metabolic vulnerability and dysregulated cerebellar signaling in somersaulting pigeons. **(A)** UMAP visualization of major cell populations identified in the cerebellum and cerebrum of homing pigeons and Xinjiang rollers. **(B)** Relative proportions of major cell types in homing pigeons and Xinjiang rollers. **(C)** Cell type–resolved expression of *GLRX2* in the cerebellum and cerebrum. Dot size indicates the fraction of cells expressing the gene; color indicates mean expression. **(D)** Cell type–resolved differences in oxidative phosphorylation (OXPHOS) UCell scores and selection-associated gene-set scores between tumbler and homing pigeons. **(E)** Pseudobulk Hallmark gene set enrichment analysis in Golgi and Purkinje cells. **(F)** Global cell–cell communication networks in the cerebellum (top) and cerebrum (bottom). **(G)** Differential incoming and outgoing signaling strengths across cerebellar cell types, highlighting Golgi and Purkinje cells as major nodes of altered communication. **(H)** Differential incoming and outgoing signaling pathways in Golgi and Purkinje cells. Shared, homing-specific, and somersaulting-specific signaling pathways are indicated. **(I)** GABA-A signaling networks in the cerebellum of homing and somersaulting pigeons. **(J)** Disease gene set enrichment across cell types, showing overlap between somersaulting-associated transcriptional changes and human brain disorder–related gene sets. **(K)** Model for *GLRX2*-linked cerebellar dysfunction in somersaulting pigeons. Reduced mitochondrial activity and stress-response dysregulation are accompanied by weakened fast inhibitory signaling and selective strengthening of excitatory and synapse-associated pathways centered on Golgi and Purkinje cells.

The cell-type composition and relative proportions were similar between somersaulting and homing pigeons in both brain regions (**Fig. 5B and table S6**), suggesting that the somersaulting phenotype does not arise from the gain or loss of specific cell populations. *GLRX2* was broadly expressed across nearly all major cell types in both regions (**Fig. 5C**) but was consistently downregulated in somersaulting pigeons. This widespread downregulation implies that the metabolic consequences of *GLRX2* loss are not restricted *a prior* to specific cell types, raising the question of whether certain populations are nonetheless preferentially affected.

To assess cell-type-specific metabolic vulnerability, we performed UCell scoring of OXPHOS gene sets. OXPHOS scores were most markedly reduced in cerebellar Golgi and Purkinje cells of somersaulting pigeons (**Fig. 5D**), whereas cerebral cell populations exhibited only modest changes, and in some cases, modestly elevated scores suggestive of compensatory response (**Fig. 5D**). Scoring with a gene set derived from loci under selection in somersaulting pigeons independently identified Golgi and Purkinje cells as the populations with the highest enrichment (**Fig. 5D**). These convergent results identify Golgi and Purkinje cells, the principle inhibitory interneurons and sole output neurons of the cerebellar cortex, respectively, with well-established roles in motor coordination and sensorimotor control (*51–53*), as the primary cellular targets in which evolutionary and metabolic signatures converge.

Pseudobulk gene set enrichment analysis (GSEA) in these two cell types revealed coordinated suppression of core mitochondrial and metabolic programs, including oxidative phosphorylation, mTORC1 signaling, reactive oxygen species pathway, glycolysis, and adipogenesis (**Fig. 5E**). Cellular stress-response and proteostatic homeostasis, the unfolded protein response, DNA repair, and protein secretion, were also downregulated (**Fig. 5E**), suggesting compromised capacity to buffer metabolic stress. Consistent with these observations, *in silico* knockout of *GLRX2* in homing-pigeon Golgi and Purkinje cells recapitulated key somersaulting-like perturbations with cell-type-specific signature that, Golgi cells showed prominent disruption of mitochondrial complex IV genes (*COX3* and *COX1*) together with axon-guidance and inhibitory synaptic programs (**fig. S16, A and C**), whereas Purkinje cells were more strongly perturbed in axon-guidance and excitatory signaling pathways (**fig. S16, B and C**). Together, these analyses reveal a shared transcriptional state in vulnerable cerebellar neurons marked by metabolic insufficiency and stress-pathway dysregulation.

We next investigated whether regional differences in metabolic vulnerability were accompanied by alterations in intercellular communication. Cell–cell communication analysis revealed that, at the global level, both the cerebellum and cerebrum of somersaulting pigeons exhibited an overall reduction in intercellular signaling, alongside a smaller subset of strengthened interactions (**Fig. 5F**). Golgi and Purkinje cells emerged as the primary nodes of altered communication in the cerebellum (**Fig. 5G**), with a consistent pattern that, fast inhibitory signaling was reduced while excitatory and synapse-associated pathways were selectively strengthen (**Fig. 5H**). Specifically, both incoming and outgoing GABA-A signaling were significantly dismissed in Golgi cells, and Purkinje cells also showed reduced incoming GABA-A signaling (**Fig. 5, H and I**). By contrast, Golgi cells exhibited increased outgoing GABA-B signaling, and increased incoming and outgoing glutamatergic, *EPHA*, *ADGRL*, and *NRXN* signaling, while Purkinje cells showed increased NRXN- and ADGRL-associated signaling (**Fig. 5H**). At the bulk level, Golgi cells showed increased incoming and outgoing signaling strength and Purkinje exhibited increased outgoing signaling strength (**Fig. 5G and fig. S17**). Signaling changes in the cerebrum were more modest and primarily involved glial rather than neuronal populations (**Fig. 5F and fig. S17**), reinforcing the regional specificity of the cerebellar circuit remodeling.

We next examined whether the dysregulated cell-type-specific programs were linked to conserved disease-associated pathways. We focused on neurological disorder gene sets related to motor control, neuronal excitability, and mitochondrial dysfunction. Epilepsy-associated gene sets were broadly enriched across both cerebellar and cerebral cell types (**Fig. 5J**), suggesting that the affected programs overlap with pathways governing neuronal excitability and circuit stability. Notably, cerebellar Purkinje cells also showed enrichment for mitochondrial disease and hereditary spastic paraplegia (HSP) gene sets (**Fig. 5J**), consistent with their mitochondrial vulnerability and central role in motor coordination. Although Golgi cells did not reach statistical significance for these disease categories, their limited representation in the dataset may have reduced the power to detect enrichment signals.

Together, these findings support a model in which *GLRX2* deficiency preferentially disrupts metabolically vulnerable cerebellar neurons, leading to remodeling fast inhibitory signaling and selective strengthening of excitatory and synapse-associated signaling in Golgi- and Purkinje-centered circuits, providing a plausible cellular and circuit-level basis for the impaired balance and aberrant somersaulting behavior.

## Discussion

This study identified dysregulation of mitochondrial redox homeostasis as a central physiological basis of the somersaulting behavior in pigeons, resolving a behavioral enigma that has persisted since Darwin’s time. Specifically, we found the loss of *GLRX2* function through an 11-bp frameshift deletion compromises mitochondrial respiratory integrity and antioxidant buffering in the brain, preferentially affecting cerebellar Golgi and Purkinje cells and remodeling the inhibitory and excitatory signaling circuits they anchor. These findings reframe the basis of this classic domesticated behavior from a neurochemical model to one centered on mitochondrial metabolic stability, and provide a rare example in which a functionally deleterious loss-of-function mutation, typically constrained by purifying selection in the wild, has been retained and enriched under artificial selection (*54*).

Although *GLRX2* loss is systemic, the behavioral phenotype implicates cerebellar circuitry. Our single-nucleus transcriptomic data offer a cellular explanation for this apparent paradox, revealing that metabolic consequences concentrate in Purkinje and Golgi cells, neurons characterized by among the substantial bioenergetic demands in the vertebrate brain (*55–57*). This selective vulnerability is consistent with the established principle that metabolically demanding neurons are particularly susceptible to OXPHOS impairment, as documented across human mitochondrial diseases from Leigh syndrome to Friedreich’s ataxia (*58–60*). These metabolic perturbations predict specific functional consequences for cerebellar circuitry. Attenuated GABA-A signaling in Golgi and Purkinje networks is consistent with degraded fast inhibitory transmission, while compensatory GABA-B upregulation and enhanced excitatory and synaptic adhesion signaling *(NRXN, ADGRL*) would not be expected to substitute for the temporal precision that ionotropic inhibition provides in motor coordination. Critically, this pathophysiology operates without detectable apoptosis, indicating chronic sublethal metabolic stress rather than neurodegeneration. This distinction helps explain the episodic and stimulus-triggered character of somersaulting, particularly in flying tumbler breeds, and aligns the phenotype with episodic ataxias and paroxysmal movement disorders rather than progressive neurodegenerative diseases (*61, 62*). However, given the broad expression and fundamental mitochondrial function of *GLRX2*, additional contributions from other cell types and tissues cannot be excluded.

The evolutionary architecture of the trait illuminates how behavioral diversity arises under artificial selection. The *GLRX2* deletion segregates at high frequency across phylogenetically distinct roller and fantail lineages, supporting a single ancestral origin, yet the broader landscape of selection signals is largely non-overlapping between lineages despite converging on shared neurobiological pathways. This pattern of a shared major-effect mutation with independent polygenic elaboration, explains how a common genetic origin has diversified into the spectrum of somersaulting phenotypes Darwin found so remarkable. The fantail case is particularly informative, as these birds carry the same deletion yet express postural instability rather than full somersaulting (*12*), demonstrating that *GLRX2* loss functions as a sensitizing mutation whose phenotypic expression depends on genetic background and selection history. Domestication enabled this trajectory by buffering the severe fitness costs of motor dysfunction, permitting the deletion’s initial persistence under relaxed purifying selection and its subsequent enrichment through artificial selection for a human-valued phenotype. Whether mitochondrial genes are disproportionately susceptible to this process, because domestication specifically relaxes selection on metabolic efficiency, is an open question that our findings motivate.

The behavioral variation among roller and tumbler breeds implies lineage-specific modifier loci that fine-tune the intensity and kinematic expression of the trait, and identifying these modifiers will clarify how a single major-effect mutation generates such phenotypic diversity. While GLRX2-deficient mice and zebrafish do not somersault, they recapitulate the conserved mitochondrial, neuropathological, and locomotor phenotypes observed in somersaulting pigeons. This consistency across three vertebrate classes strengthens the case for *GLRX2* as a major contributor to the somersaulting phenotype, though definitive demonstration of its sufficiency will require gene editing in pigeons, a technical frontier that remains challenging but is advancing rapidly. More broadly, cell-type-specific perturbation and electrophysiological approaches in avian models will be needed to resolve how a core mitochondrial defect translates into the diverse motor phenotypes that have fascinated biologists since Darwin’s time.

## Acknowledgments

We thank Leif Andersson (Uppsala University) and Nicolas Alexandre (Colossal Biosciences) for their helpful discussions and comments. We thank the Animal Bank at the Germplasm Bank of Wild Species (https://cstr.cn/31121.02.GBOWS.AB) for providing biological materials and technical support. We also thank the Institutional Center for Shared Technologies and Facilities of Kunming Institute of Zoology (KIZ), Chinese Academy of Sciences (CAS), for our Electron Microscopy, Yingqi Guo, Xingcai Wu, and Rongrong Fu for their help of making EM sample and taking EM images, Yuchen Gong for sharing her expertise and providing guidance on zebrafish behavioral experiments, as well as Linyao Du and Menghan Qiu for filming the flight video of Xinjiang rollers. Photo, video production and sampling were supported by the University of Debrecen Program for Scientific Publication.

## Funding

This research was supported by

National Science Foundation of China grant 32400353 (SS)

Yunnan Fundamental Research Projects 202301AT070314 (SS)

National Science Foundation of China grant 32470453 (MSW)

Yunnan Fundamental Research Projects 202401AV070007 (MSW)

Yunnan Fundamental Research Projects 202305AH340006 (MSW)

Iran National Science Foundation 032875 (AE)

## Author contributions

Conceptualization: SS, MSW

Methodology: SS, LYS, MLL

Investigation: SS, LYS, LL, YQ, WDM, XG

Validation: LYS, WDM, CZ

Formal analysis: SS, KJ, GZZ

Resources: MSW, AE, SK, JL, LM, LYS, QP, WJP, CRL, JL, ZB, DB, HKK, JP, JLD

Data curation: SS

Funding acquisition: SS, MSW, AE

Visualization: SS, HMC, SK, ZB, GHW

Project administration: SS, MSW

Supervision: MSW

Writing – original draft: GZ, MSW, SS

Writing – review & editing: SS, MSW, YGY, LYS, ZW, LG, CY, LZL, ZL, ZB, WG

## Competing interests

MSW, CZ, SS, WDM, and LL declare that they have filed a patent application relating to the GLRX2 11-bp mutation associated with the tumbling phenotype in pigeons. All other authors declare that they have no competing interests.

## Data, code, and materials availability

The raw and processed data generated in this study have been deposited in the Genome Sequence Archive (GSA) under accession number CRA015615.

## Supplementary Materials

Materials and Methods

Figs. S1 to S17

Tables S1 to S8

References (63–113)

Movies S1 to S5

